# Real-time near-infrared imaging distinguishes nasopharyngeal colonization from aspiration of *Streptococcus pneumoniae* and identifies aspiration as a trigger of severe disease

**DOI:** 10.64898/2026.08.21.746383

**Authors:** Takashi Saito, Momoko Kobayashi, Zhengkuan Sun, Satoshi Muraoka, Daisuke Motooka, Takako Yoshida, Shin-ichi Shiomi, Jun Adachi, Masaya Yamaguchi

## Abstract

*Streptococcus pneumoniae* asymptomatically colonizes the nasopharynx but can invade the lower respiratory tract to cause life-threatening disease, particularly in older adults. However, whether the initial site of bacterial deposition following intranasal inoculation determines disease progression has not been directly examined. Here, we developed a near-infrared (NIR) fluorescence imaging approach using indocyanine green (ICG)-labeled *S. pneumoniae* TIGR4 to visualize early bacterial distribution in real time. ICG labeling by simple mixing, without genetic or chemical modification, neither impaired bacterial growth at 33 or 37 °C, nor altered acid tolerance. Continuous video imaging during the first 10 min of infection resolved two distinct patterns: bacteria confined to the nasopharynx (colonization) and those aspirated into the lower respiratory tract (aspiration). Kaplan–Meier analysis revealed markedly higher mortality in the aspiration group in both young (hazard ratio = 7.9) and aged (hazard ratio = 8.4) male C57BL/6J mice, despite a 10-fold lower inoculum used for aged animals, with deaths beginning on day 3. Systemic profiling of blood at 24 h by RNA sequencing and plasma proteomics revealed that early aspiration in aged mice was associated with the activation of inflammatory and hematopoietic programs, enrichment of complement and coagulation cascades, and phagocytic pathways. Together, these findings establish aspiration into the lower respiratory tract as a trigger of severe pneumococcal disease and introduce real-time NIR imaging as a technique for linking early infection dynamics to systemic host responses.

## Introduction

*Streptococcus pneumoniae* asymptomatically colonizes the human nasopharynx and is carried by 27%–65% of healthy children, compared to < 10% of adults, identifying children as a major reservoir [1]. Despite this commensal relationship, the bacterium can invade sites it does not normally colonize, including the lower respiratory tract, bloodstream, and central nervous system, thereby causing pneumonia, bacteremia, and meningitis [1, 2]. The global burden of pneumococcal disease remains substantial: *S. pneumoniae* has been estimated to account for approximately 829,000 deaths worldwide in 2019 [3], with disproportionately high morbidity and mortality in young children, older adults [4], and immunocompromised individuals [5]. Conjugate vaccines, such as PCV10 and PCV13, reduce vaccine-serotype diseases; however, serotype replacement and persistence of certain serotypes result in a continued and considerable burden of pneumococcal disease, motivating the development of higher-valency and protein-based vaccines [5, 6].

Older age is a major risk factor for invasive pneumococcal disease and is associated with poor outcomes [7]. Two interrelated processes, immunosenescence and inflammaging, probably underlie this vulnerability: immunosenescence refers to age-associated decline in immune function, whereas inflammaging describes a chronic low-grade inflammatory state, which accumulates with age [8, 9]. Studies using aged mouse models have identified several contributing mechanisms: innate immune deficits prolong nasopharyngeal carriage [7]; tumor necrosis factor-mediated monocyte dysfunction blunts antibacterial responses [10]; and excessive inflammation coupled with diminished interleukin-10 production heightens disease susceptibility [11]. Furthermore, inflammaging-associated cytokines may upregulate receptors, such as platelet-activating factor receptor on host cells, thereby facilitating bacterial adhesion and invasion [12, 13]. Collectively, these findings implicate age-related immune dysregulation, rather than a single pathway, in heightened susceptibility of older hosts. In particular, age-related decline in swallowing function makes aspiration an important route, through which nasopharyngeal bacteria reach the lower respiratory tract in older adults, and aspiration pneumonia is a major cause of morbidity and mortality in this population [14]. However, how early bacterial localization and dissemination shape subsequent disease progression remains unclear [15, 16].

Indocyanine green (ICG) is a clinically approved near-infrared (NIR) fluorescent dye with excitation and emission wavelengths of approximately 780 and 830 nm, respectively, and a well-characterized clinical safety profile [17]. Because NIR light penetrates biological tissues more effectively than does visible light, and endogenous autofluorescence is minimal in this spectral range, ICG imaging achieves high signal-to-background ratios and can detect bacteria even after they spread into the lower respiratory tract. ICG-based agents have been investigated for imaging bacterial infection and antibacterial photodynamic or photothermal therapy [18, 19]. However, resolving the spatial distribution of bacteria during the first minutes after intranasal inoculation requires continuous, real-time acquisition, which conventional bioluminescence imaging cannot provide because it captures images only sequentially. Therefore, we used a dual-sensor NIR imaging system that can record visible and fluorescence images simultaneously, enabling continuous video acquisition of bacterial distribution in live animals and unambiguous distinction between animals with signals confined to the nasopharynx and those in which the signal extends into the lower airway.

Pneumococcal pathogenesis has been extensively studied; however, whether the initial site of bacterial deposition following intranasal inoculation determines disease progression has not been directly examined. In the present study, we developed an ICG-labeling approach to visualize S. pneumoniae in real time following intranasal infection in mice. Aspiration has long been recognized as a primary route of pneumococcal invasion into the lower respiratory tract [1, 2]; however, we directly visualized this process as it occurred and linked early distribution patterns to subsequent survival outcomes. Furthermore, to investigate the potential effect of host aging on the dynamics of pneumococcal infection, we compared bacterial dissemination patterns and disease progression between young and aged mice. We further profiled the systemic host response at 24 h after inoculation to explore how early aspiration is reflected in circulating proteome and transcriptome.

## Materials and methods

### Materials

#### Reagents and chemicals

Indocyanine green (ICG) was purchased from Tokyo Chemical Industry Co., Ltd. (Tokyo, Japan). A depilatory cream (Epilat Sensitive) was from Kracie Home Products, Ltd. (Tokyo, Japan). For proteomic sample preparation: tris(2-carboxyethyl)phosphine hydrochloride (#209-19861; Fujifilm Wako Pure Chemical Corporation, Osaka, Japan), iodoacetamide (#19302-54; Nacalai Tesque, Inc., Kyoto, Japan), Trypsin/Lys-C (Promega, Madison, WI, USA), and ReproSil-Pur C18-AQ 1.9 µm resin (Dr. Maisch GmbH, Ammerbuch, Germany) were used.

#### Kits

NucleoSpin RNA Blood kit (Takara Bio Inc., Shiga, Japan); TruSeq Stranded mRNA Library Prep Kit (Illumina, Inc., San Diego, CA, USA); MagCapture Exosome Isolation HTS kit (#293-96401; Fujifilm Wako Pure Chemical Corporation, Osaka, Japan); SP3 paramagnetic particles, hydrophilic (#45152105050250) and hydrophobic (#65152105050250) (Thermo Fisher Scientific, Waltham, MA, USA).

#### Anesthetics and related drugs

Anesthesia was induced with a combination of three injectable agents—medetomidine hydrochloride (Domitor; Nippon Zenyaku Kogyo Co., Ltd., Fukushima, Japan), midazolam (Dormicum; Maruishi Pharmaceutical Co., Ltd., Osaka, Japan), and butorphanol tartrate (Vetorphale; Meiji Seika Pharma Co., Ltd., Tokyo, Japan). Atipamezole hydrochloride (Antisedan; Nippon Zenyaku Kogyo Co., Ltd., Fukushima, Japan), an α2-adrenergic antagonist, was administered to accelerate recovery from anesthesia. Isoflurane (Viatris Healthcare G.K., Tokyo, Japan) was used for inhalational general anesthesia.

#### Instruments

LuminousQuester NX fluorescence imaging system (Shimadzu Corporation, Kyoto, Japan), equipped with an external excitation laser (760 nm; total output, 1,010 mW over a 5-cm-diameter circular field, corresponding to an irradiance of approximately 51.4 mW/cm²) and an 830 nm emission filter; VICTOR Nivo multimode microplate reader (Revvity, Waltham, MA, USA); NovaSeq X Plus sequencing platform (Illumina, Inc., San Diego, CA, USA); Agilent 2100 Bioanalyzer (Agilent Technologies, Santa Clara, CA, USA); Qubit fluorometer (Thermo Fisher Scientific, Waltham, MA, USA); LTQ-Orbitrap Astral mass spectrometer coupled to a Vanquish Neo UHPLC (Thermo Fisher Scientific, Waltham, MA, USA); KingFisher Apex system (Thermo Fisher Scientific, Waltham, MA, USA).

#### Ethics statement

All animal experiments were approved by the Institutional Animal Care and Use Committee of National Institutes of Biomedical Innovation, Health and Nutrition (approval number: DSR06-24). For intranasal inoculation and imaging, mice were anesthetized by isoflurane followed by intraperitoneal injection of a three-agent combination anesthetic (medetomidine hydrochloride, 0.6 mg/kg; midazolam, 8.0 mg/kg; and butorphanol tartrate, 10.0 mg/kg). Atipamezole hydrochloride (6.0 mg/kg) was intraperitoneally administered after imaging to accelerate anesthetic recovery. At the end of experiments, mice were euthanized by cervical dislocation. In vivo experiments were performed between September 2025 and April 2026, and blood sampling with subsequent RNA-seq and plasma proteomic analyses was performed between December 2025 and July 2026.

#### Preparation of ICG–SPT4

SPT4, a clinical isolate, was cultured in Todd-Hewitt broth, supplemented with 0.2% yeast extract (THY medium) as previously described [20, 21].

SPT4 was cultured in THY medium at 37 °C, harvested by centrifugation (5,800 ×g, 10 min, 4 °C), and resuspended in phosphate-buffered saline (PBS). The suspension was adjusted so that each mouse received a 20-µL intranasal inoculum containing 1 × 10⁷ or 1 × 10⁶ colony-forming units (CFU) for young or aged mice, respectively. ICG was then added to each inoculum to a final concentration of 10 µg/mL and incubated at room temperature for 1 min; unbound ICG was removed by centrifugation (5,800 ×g, 10 min, 4 °C), and the pellet was resuspended in PBS to yield the final inoculum.

#### Growth characteristics of ICG–SPT4

To assess whether ICG labeling affected bacterial growth, the optical density at 600 nm (OD₆₀₀) was monitored over 24 h using a VICTOR Nivo multimode microplate reader. SPT4 and ICG–SPT4 inocula were serially diluted 10-fold 4 times in PBS. The diluted suspensions were seeded in 96-well plates (200 µL/well) and incubated at 33 or 37 °C, with OD₆₀₀ measured at 30-min intervals. For each well, the maximum specific growth rate (µ_max_) was calculated as the maximum slope of natural-log-transformed OD₆₀₀ over a sliding window, and total growth was quantified as the area under the OD₆₀₀–time curve (0–24 h) following the trapezoidal rule (n = 6 wells per group). For each temperature and metric, the four groups (unlabeled and ICG-labeled bacteria at each of the two inoculum concentrations) were compared using ordinary one-way analysis of variance followed by Šídák’s multiple-comparisons test. Homogeneity of variance was confirmed using the Brown–Forsythe and Bartlett tests, and normality of residuals using the Shapiro–Wilk test.

#### Real-time NIR fluorescence imaging of intranasal bacterial inoculation in vivo

Male C57BL/6J mice aged 2 months (2M; young) and 18 months (18M; aged) were used in this study. Male mice were used exclusively to minimize potential confounding effects of hormonal variation on immune responses and infection outcomes [22]. Ventral fur was removed using a depilatory cream (Epilat Sensitive) prior to imaging. Young mice were obtained as C57BL/6J (CLEA Japan, Inc., Tokyo, Japan), and aged mice were obtained as C57BL/6J (RIKEN BioResource Research Center, Tsukuba, Japan) through the National BioResource Project of MEXT/AMED, Japan. Mice were housed under specific-pathogen-free (SPF) conditions at 24–26 °C and 50–60% relative humidity under a 12-h light/dark cycle (lights on at 07:00), with free access to a γ-irradiated (10 kGy) standard diet (FF; Funabashi Farm Co., Ltd., Chiba, Japan) and water, and were acclimatized for 7–14 days before the experiments. ICG–SPT4 was intranasally administered as four doses of 5 µL per nostril at 2-min intervals (20 µL total). Bacterial distribution was monitored for 10 min using a LuminousQuester NX fluorescence imaging system (760 nm excitation laser, irradiance ≈ 51.4 mW/cm², 830 nm emission filter). Mice with fluorescence confined to the nasopharynx were classified as colonization, and those with signal extending into the lower respiratory tract as aspiration. Survival was monitored daily for 14 days. Animals that reached humane endpoints were euthanized and recorded as deaths. Survival was estimated by the Kaplan–Meier method, and survival curves of colonization and aspiration groups were compared using the log-rank (Mantel–Cox) test, with hazard ratios calculated by the same method. Survival analyses were performed using GraphPad Prism v.9.5.1 (GraphPad Software, San Diego, CA, USA), and *P* < 0.05 was considered statistically significant.

#### Mouse infection and sample collection for omics analyses

Mouse infection assays were performed as previously described [20, 23]. Young (2M) and aged (18M) mice were intranasally inoculated with SPT4. At 24 h post-inoculation, cardiac blood was collected, and plasma was isolated. Mice were divided into six groups based on age and infection route: 2M control (Young_Ctrl, n = 6), 18M control (Aged_Ctrl, n = 3), 2M colonization (Young_Col, n = 2), 2M aspiration (Young_Asp, n = 4), 18M colonization (Aged_Col, n = 4), and 18M aspiration (Aged_Asp, n = 4).

#### Simulated gastric fluid (SGF) assay

SPT4 and ICG–SPT4 were prepared as described above. SGF was prepared according to the US Pharmacopeia formulation [24], by dissolving NaCl (2 mg/mL) and pepsin (3.2 mg/mL) in Milli-Q water, with pH adjusted to 1.0–1.2 (fasting) using 1 N HCl or 3.8–4.1 (postprandial) without HCl. Bacterial survival in SGF was assessed as previously described [25]. The 10-fold concentrated suspension was added to 1 mL SGF (20 µL per mL) and incubated at 37 °C. At 5, 15, 30, and 60 min, 100-µL aliquots were withdrawn, serially diluted 10-fold 3 times in PBS, and plated in triplicate (20 µL per well) on blood agar. Colonies were counted after overnight incubation at 37 °C.

#### Bulk RNA sequencing (RNA-seq)

RNA-seq of mouse blood was performed as previously described [26]. Cardiac blood was collected 24 h post-inoculation, and RNA was extracted using a NucleoSpin RNA Blood kit according to the manufacturer’s instructions. RNA integrity was confirmed using an Agilent 2100 Bioanalyzer (RNA integrity number ≥ 7.5 for all samples), and RNA concentration was measured using a Qubit fluorometer. Libraries were prepared using a TruSeq Stranded mRNA Library Prep Kit and sequenced on a NovaSeq X Plus platform in 2 × 151-bp paired-end mode. Two samples per group were analyzed: Young_Col, Young_Asp, Aged_Col, and Aged_Asp. After adapter trimming and alignment to the mouse reference genome (GRCm39, GENCODE v.M35) using STAR, each library yielded 61 million paired-end reads (range, 55.6–65.0 million), of which 40%–48% were uniquely mapped to the reference genome, a relatively low rate attributable to abundant globin and ribosomal RNA transcripts in whole-blood samples.

#### Read processing and differential expression analysis

Raw read quality was assessed using FastQC v.0.11.9 and MultiQC v.1.35 [27]. Adapters were trimmed using fastp v.1.1.0 with automatic detection for paired-end reads [28]. Trimmed reads were aligned to the mouse reference genome (GRCm39) using STAR v.2.7.10a [29] with the GENCODE v.M35 annotation [30]. Gene-level counts were obtained using the --quantMode GeneCounts option (strand2 column, selected by column-sum comparison confirming library orientation) in STAR, and Ensembl gene IDs were mapped to symbols using the GENCODE v.M35 GTF file.

Differentially expressed genes (DEGs) were identified using DESeq2 [31] as implemented in pyDESeq2 v.0.5.4 [32]; genes with nominal P < 0.05 and |log_2_FC| > 1 were defined as DEGs. Because of the small sample size (n = 2 per group), adjusted P-values were not used as the primary threshold. Four pairwise comparisons were performed: Young_Asp vs. Young_Col, Aged_Asp vs. Aged_Col, Aged_Asp vs. Young_Asp, and Aged_Col vs. Young_Col.

#### Visualization of transcriptomic data and pathway analysis

Principal component analysis (PCA) was performed on z-score-normalized, log_2_-transformed, size-factor-normalized counts using scikit-learn v.1.8.0 [33]. Volcano plots and hierarchical clustering heatmaps (average linkage, Euclidean distance; 50 top DEGs ranked by P-value) were generated using matplotlib v.3.10.8 [34] and seaborn v.0.13.2 [35]. Gene set enrichment analysis (GSEA) was performed on a preranked list (ranked by sign(log_2_FC) × (-log₁₀(*P-*value))) using the prerank module of gseapy v.1.1.13 [36] with 100 permutations and a fixed random seed of 42, against the Kyoto Encyclopedia of Genes and Genomes (KEGG) 2019 Mouse and Gene Ontology (GO) Biological Process 2023 gene set libraries obtained from Enrichr [37, 38], with pathways at false discovery rate (FDR) q-value < 0.25 considered significant. GSEA was used for the transcriptomic data because the full ranked gene list was available, whereas over-representation analysis was used for the proteomic data, which were based on discrete differentially abundant protein lists. For visualization, the top 10 significant pathways ranked by FDR q-value are shown.

#### Sample preparation

Label-free proteomic analysis of plasma-derived extracellular vesicles (EVs) was performed as previously described [39]. Briefly, EVs were isolated from heparinized plasma using a MagCapture Exosome isolation HTS kit and KingFisher Apex system. Isolated EVs were lysed using a phase-transfer surfactant buffer, and EV proteins were reduced with 10 mM tris(2-carboxyethyl)phosphine hydrochloride, alkylated with iodoacetamide, and processed using an automated single-pot, solid phase-enhanced sample-preparation (SP3) workflow [40] on a KingFisher Apex system. Proteins were digested with Trypsin/Lys-C, and the resulting peptides were acidified, desalted using C18-SCX StageTips, and subjected to liquid chromatography–tandem mass spectrometry (LC–MS/MS) analysis [41].

#### LC–MS/MS analysis

Nano-LC–MS/MS analysis was performed on an LTQ-Orbitrap Astral mass spectrometer coupled to a Vanquish Neo UHPLC system. Peptides were separated on an in-house-packed analytical column (75 µm × 30 cm, ReproSil-Pur C18-AQ, 1.9 µm resin) over a 9.5-min gradient of acetonitrile in 0.1% formic acid, followed by high-organic wash and re-equilibration (16 min total per run). Data were acquired in data-independent acquisition mode.

#### DIA LC–MS/MS acquisition parameters

Full MS scans were recorded in the Orbitrap over *m/z* 380–980 at a resolution of 240,000, with an AGC target of 5 × 10⁶ and a maximum injection time of 5 ms. DIA MS/MS spectra were acquired in the Astral analyzer using 300 variable 2-*m/z* isolation windows spanning m/z 380–980; fragment ions were detected over *m/z* 150–2000 with an AGC target of 5 × 10⁴, a maximum injection time of 3 ms, and a normalized collision energy of 25%. All data were acquired in positive-ion mode in profile format. The gradient was: solvent B (99.9% acetonitrile, 0.1% formic acid [FA]) raised from 5% to 25% over 4 min and from 25% to 50% over the next 5.5 min; followed by a high-organic wash and re-equilibration (16 min total per run); solvent A was 0.1% FA in water.

#### Database search and protein quantification

Raw MS data were processed using the DIA-NN software v.2.2.0 [42]. Spectral library-free database searching was performed against the UniProt Mus musculus reference proteome database (taxonomy ID: 10090; downloaded in May 2025), supplemented with a contaminant database. The search parameters were set as follows: a maximum of two missed cleavages, peptide lengths ranging from 7 to 30 amino acids, and carbamidomethylation of cysteine residues (+57.021 Da) as a fixed modification. Quantification was performed using the QuantUMS strategy, and RT-dependent normalization was used for cross-run normalization. Peptide precursor identifications were filtered at 1% FDR.

#### Data filtering and imputation of missing values

Proteins detected in at least 70% of total samples (≥16 of 23 samples) were retained for downstream analysis, resulting in 4,006 proteins. Missing values were imputed using the within-group minimum value; if all values within a group were missing, the global minimum across all samples was used instead.

#### Differential abundance analysis

Differentially expressed proteins (DEPs) were identified using Welch’s t-test implemented in SciPy v.1.17.1 [43]. Proteins with P < 0.05 and |log_2_FC| > 0.5 were defined as DEPs. The following three comparisons were performed as primary analyses: Aged_Ctrl vs. Young_Ctrl, Aged_Asp vs. Young_Asp, and Aged_Asp vs. Aged_Col. Two additional comparisons, Young_Asp vs. Young_Col and Aged_Col vs. Young_Col, were performed as supplementary analyses and interpreted with caution because of the limited sample size of the Young_Col group (n = 2).

#### Visualization of proteomic data and pathway analysis

PCA was conducted, and volcano plots and hierarchical clustering heatmaps (average linkage, Euclidean distance) were generated, as described for the transcriptomic data. Protein–protein interaction (PPI) networks were constructed using the STRING database v.12.0 (*M. musculus*, confidence score ≥ 0.4) [44] and visualized using networkx v.3.6.1 [45]. Over-representation analysis was performed using the enrichr module of gseapy v.1.1.13 [36] against the KEGG 2019 Mouse and GO Biological Process 2023 gene set libraries obtained from Enrichr [37, 38], with Benjamini–Hochberg-adjusted *P* < 0.05 considered significant. Representative KEGG pathway diagrams were visualized using KEGG Mapper (accessed May 2026) [46, 47]. For visualization, the top 15 significant pathways ranked by nominal P-value are shown.

#### Multi-omics integration

To identify genes with concordant transcript- and protein-level changes, RNA-seq and proteomic datasets were integrated for four matched comparisons: Young_Asp vs. Young_Col, Aged_Asp vs. Aged_Col, Aged_Asp vs. Young_Asp, and Aged_Col vs. Young_Col. Datasets were merged on gene symbol, and the overlap between significant DEGs (|log_2_FC| > 1, *P* < 0.05) and DEPs (|log_2_FC| > 0.5, *P* < 0.05) was visualized as Venn diagrams. Transcript- and protein-level log_2_FC values were compared by Pearson’s correlation for all genes detected in both datasets.

## Results

### ICG labeling does not impair the growth kinetics of S. pneumoniae strain TIGR4

*S. pneumoniae* strain TIGR4 (SPT4) was labeled with ICG (ICG–SPT4). The growth kinetics of ICG–SPT4 was indistinguishable from that of unlabeled SPT4. The growth curves of SPT4 and ICG–SPT4 overlapped throughout the 24-h culture period at both 33 and 37 °C, and young- and aged-mouse inoculum concentrations (Figure 1A, B). The µ_max_ and total growth (the area under the OD₆₀₀–time curve over 0–24 h) did not significantly differ at either temperature when SPT4 and ICG–SPT4 were compared within each inoculum concentration (Figure 1C, D), indicating that ICG labeling did not affect the growth of SPT4.

**Figure 1.**
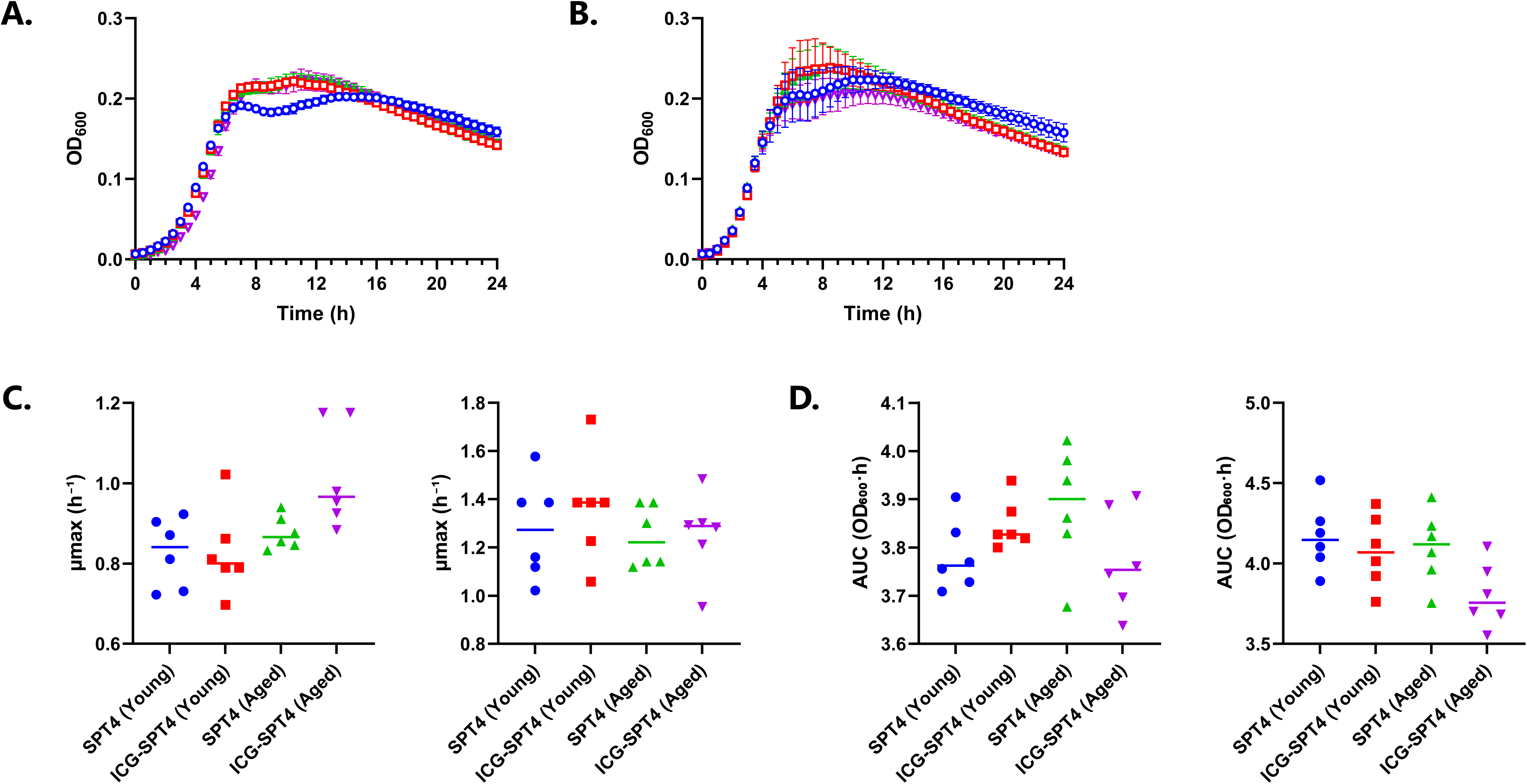
Growth characteristics of ICG–SPT4. (A) Growth curves of SPT4 and ICG–SPT4 at 33 °C, simulating nasopharyngeal temperature conditions. (B) Growth curves at 37 °C, simulating systemic temperature conditions. In (A) and (B), each strain was prepared at the young- and aged-mouse inoculum concentrations: SPT4 at young (blue circles) and aged (green upward triangles) concentrations, and ICG–SPT4 at young (red squares) and aged (purple downward triangles) concentrations. Bacteria were serially diluted 10-fold 4 times in PBS prior to seeding in 96-well plates (200 µL/well). OD₆₀₀ was measured at 30-min intervals for 24 h. Data are presented as mean ± standard deviation (n = 6 wells per group). (C) Maximum specific growth rate (µ_max_) and (D) total growth (area under the OD₆₀₀–time curve; AUC, 0–24 h) at 33 °C (left) and 37 °C (right). Each symbol represents an individual well, and the horizontal bar indicates the group mean (n = 6 wells per group). SPT4 and ICG–SPT4 were compared within each inoculum concentration using ordinary one-way analysis of variance with Šídák’s multiple-comparisons test.

### Real-time NIR fluorescence imaging captures two distinct bacterial distribution patterns in vivo

To visualize early bacterial distribution following intranasal inoculation, ICG–SPT4 was administered to aged mice and tracked by real-time NIR fluorescence imaging. Representative images from colonization (Figure 2A) and aspiration (Figure 2B) mice illustrate two distinct distribution patterns observed within the first 10 min post-inoculation. In the colonization mouse, NIR fluorescence signal remained confined to the nasopharyngeal region throughout the observation period (Figure 2A). In contrast, in the aspiration mouse, the signal was first detected extending into the lower respiratory tract 4 min post-inoculation and persisted through the end of the observation period (Figure 2B).

**Figure 2.**
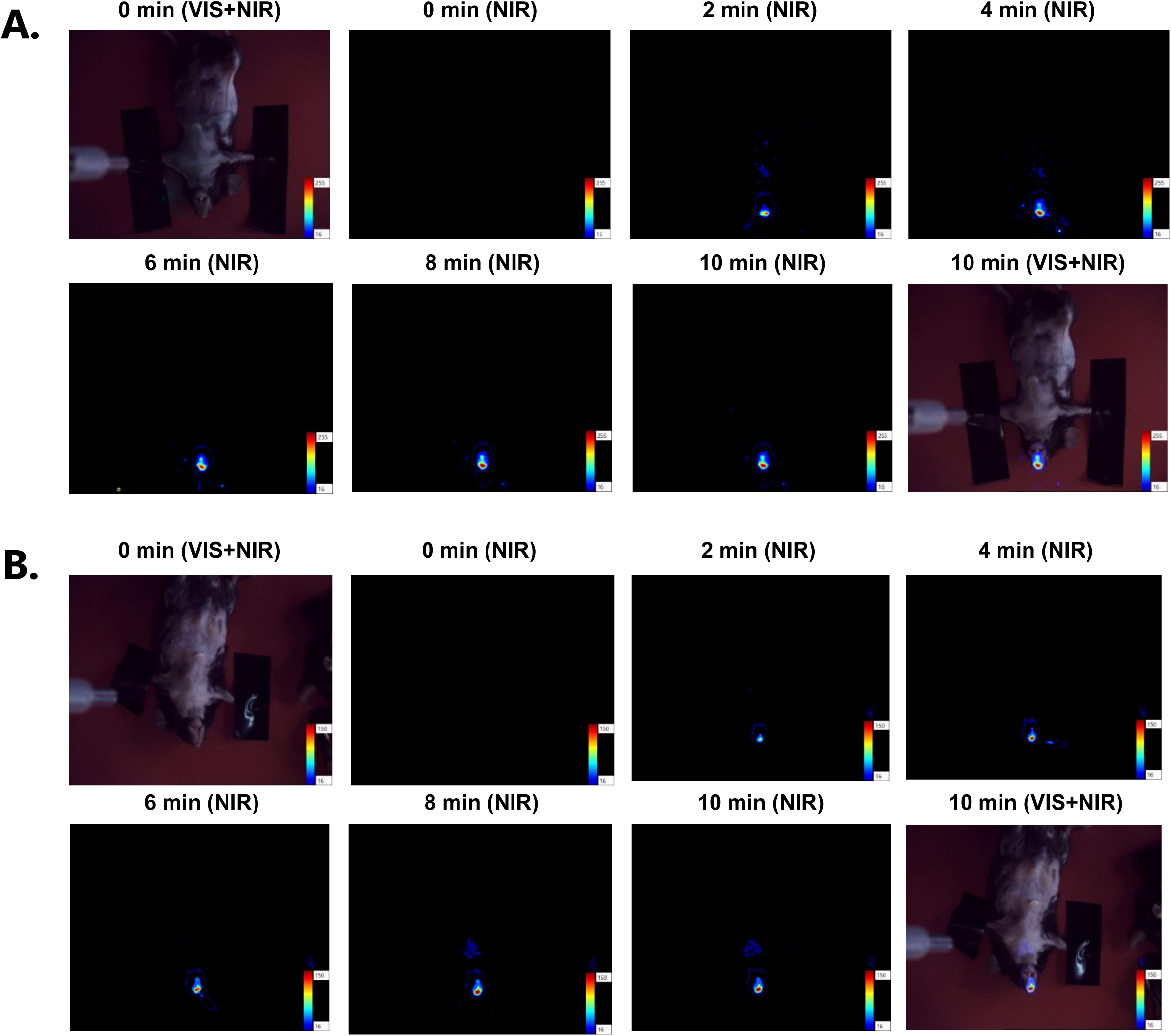
Real-time NIR fluorescence imaging of ICG–SPT4 following intranasal inoculation. Representative sequential images from aged colonization (A) and aspiration (B) mice. Images were acquired before inoculation (0 min; visible (VIS) +NIR overlay), immediately after the first inoculation (0 min; NIR) and at 2-min intervals following each subsequent dose (2, 4, 6, 8 and 10 min; NIR), with the final image at 10 min post-inoculation (VIS+NIR overlay). ICG–SPT4 was intranasally administered in four doses at 2-min intervals. Mice were assigned to the colonization or aspiration group based on the presence or absence of lower respiratory tract signal within 10 min of inoculation. Color scale indicates fluorescence intensity (arbitrary units). See Supplementary Videos 1 and 2 (https://doi.org/10.6084/m9.figshare.33457540) for representative Aged_Asp time-lapse recordings.

### Continuous real-time acquisition, unlike conventional still-frame optical imaging, reliably captures the transient aspiration signal

Real-time whole-body tracking of ICG–SPT4 during the first minutes after intranasal inoculation was performed with a benchtop NIR fluorescence imaging system (LuminousQuester NX, Shimadzu). This system simultaneously captures a visible-light image and an NIR fluorescence image and overlays them in real time at up to 10 frames per second, providing continuous video of bacterial distribution. Because it relies on NIR light, imaging is performed on an open benchtop under room lighting (free of near-infrared wavelengths) without a light-tight enclosure, so that animals can be positioned and dosed during acquisition. An external 760 nm excitation laser was used, as an excitation light source is not supplied with the instrument.

In contrast, conventional in vivo optical imagers such as the IVIS platform (Revvity/PerkinElmer) operate inside a light-tight cabinet and acquire sequential still images with a cooled CCD, using exposure times on the order of seconds to minutes and switching excitation/emission filters between acquisitions. Although these systems support kinetic acquisition of consecutive still frames, they do not provide continuous real-time video of the seconds-to-minutes distribution phase that is central to the present study.

Conventional *in vivo* optical imaging distinguished colonization- and aspiration-type distributions on individual still images (Figure S3) but could not reliably classify the two. Because still-frame acquisition requires an enclosed cabinet and sequential mechanical and optical operations, imaging was limited to a single time point after inoculation and could not follow the distribution phase in real time. The aspirated signal in the lower airway was only transient: on serial acquisitions it faded and was ultimately lost (data not shown). Owing to its low abundance and rapid dispersion, this signal did not persist long enough to form a detectable focus during the long integrating exposure required by such systems—a limitation reflecting the dynamic nature of the signal rather than insufficient sensitivity, as these systems detect weak but spatially stationary signals well. As a result, true aspiration events could be missed and misclassified as colonization, contaminating the colonization group. Continuous real-time imaging avoids this misclassification by capturing the transient aspiration signal throughout the inoculation sequence, allowing accurate binary assignment.

### Aspiration into the lower respiratory tract markedly increases mortality across host ages

Kaplan–Meier analysis revealed a significant difference in survival between the colonization and aspiration groups in both age cohorts (Figure 3). In young mice inoculated with 1.0–1.7 × 10⁷ CFU/mouse, survival was markedly higher in the colonization group than in the aspiration group (hazard ratio = 7.9, 95% CI 1.5–40.6, log-rank test *P* = 0.022; Figure 3A). A comparable pattern was observed in aged mice inoculated with 1.1–1.9 × 10⁶ CFU/mouse (hazard ratio = 8.4, 95% CI 2.4–29.1, log-rank test *P* = 0.013; Figure 3B). These results demonstrate that early aspiration into the lower respiratory tract is associated with substantially increased mortality across host ages, despite a 10-fold difference in inoculum doses between the two cohorts.

**Figure 3.**
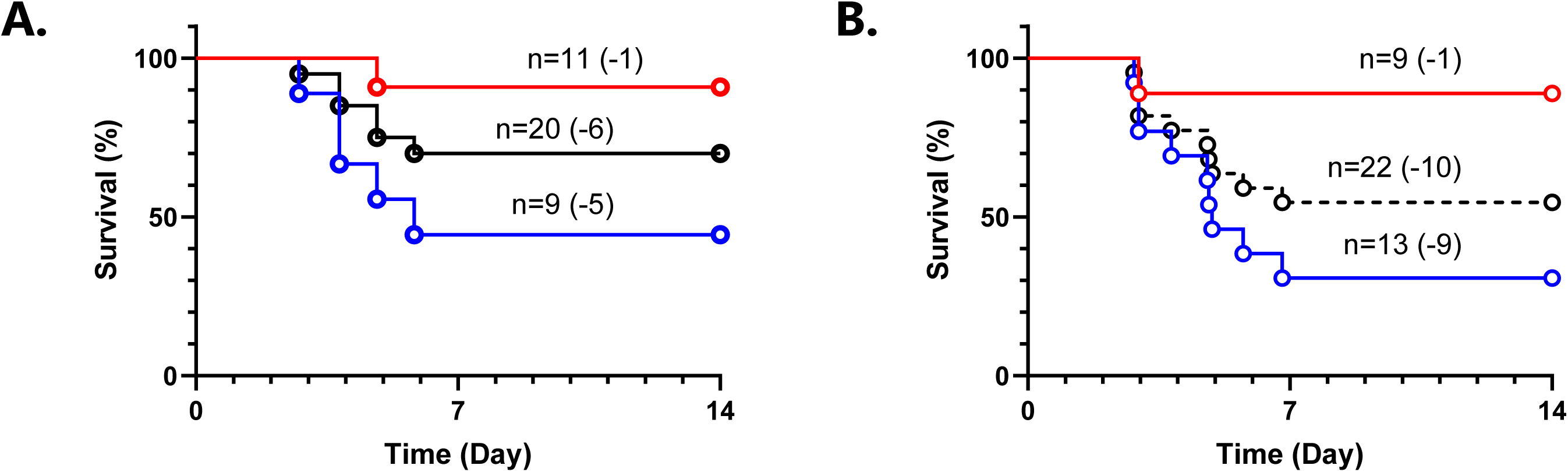
Survival analysis following intranasal inoculation of ICG–SPT4. Kaplan–Meier survival curves for young mice inoculated with 1.0–1.7 × 10⁷ CFU/mouse (A) and aged mice inoculated with 1.1–1.9 × 10⁶ CFU/mouse (B). Mice were stratified into colonization (Col, red) and aspiration (Asp, blue) groups based on NIR fluorescence imaging within the first 10 min post-inoculation. Total (black dashed) indicates all inoculated mice regardless of group assignment. Numbers in the plots indicate the group size, with the number of deaths during the 14-day observation period in parentheses (young: Col, n = 11 [1 death]; Asp, n = 9 [5 deaths]; total, n = 20 [6 deaths]. Aged: Col, n = 9 [1 death]; Asp, n = 13 [9 deaths]; total, n = 22 [10 deaths]).

### SPT4 is rapidly killed under simulated gastric conditions

To assess whether ICG–SPT4 could survive gastric conditions, we exposed SPT4 and ICG–SPT4 to SGF at pH 1.0–1.2 (fasting) or 3.8–4.1 (postprandial) for up to 60 min (Figure 4). At pH 1.0–1.2, viable bacteria were undetectable within 5 min in both SPT4 and ICG–SPT4; at pH 3.8–4.1, viability was completely lost within 15 min in both groups. In contrast, bacteria suspended in PBS retained viability throughout the observation period. SPT4 and ICG–SPT4 showed comparable susceptibility profiles under all conditions, further indicating that ICG labeling did not substantially alter bacterial physiology.

**Figure 4.**
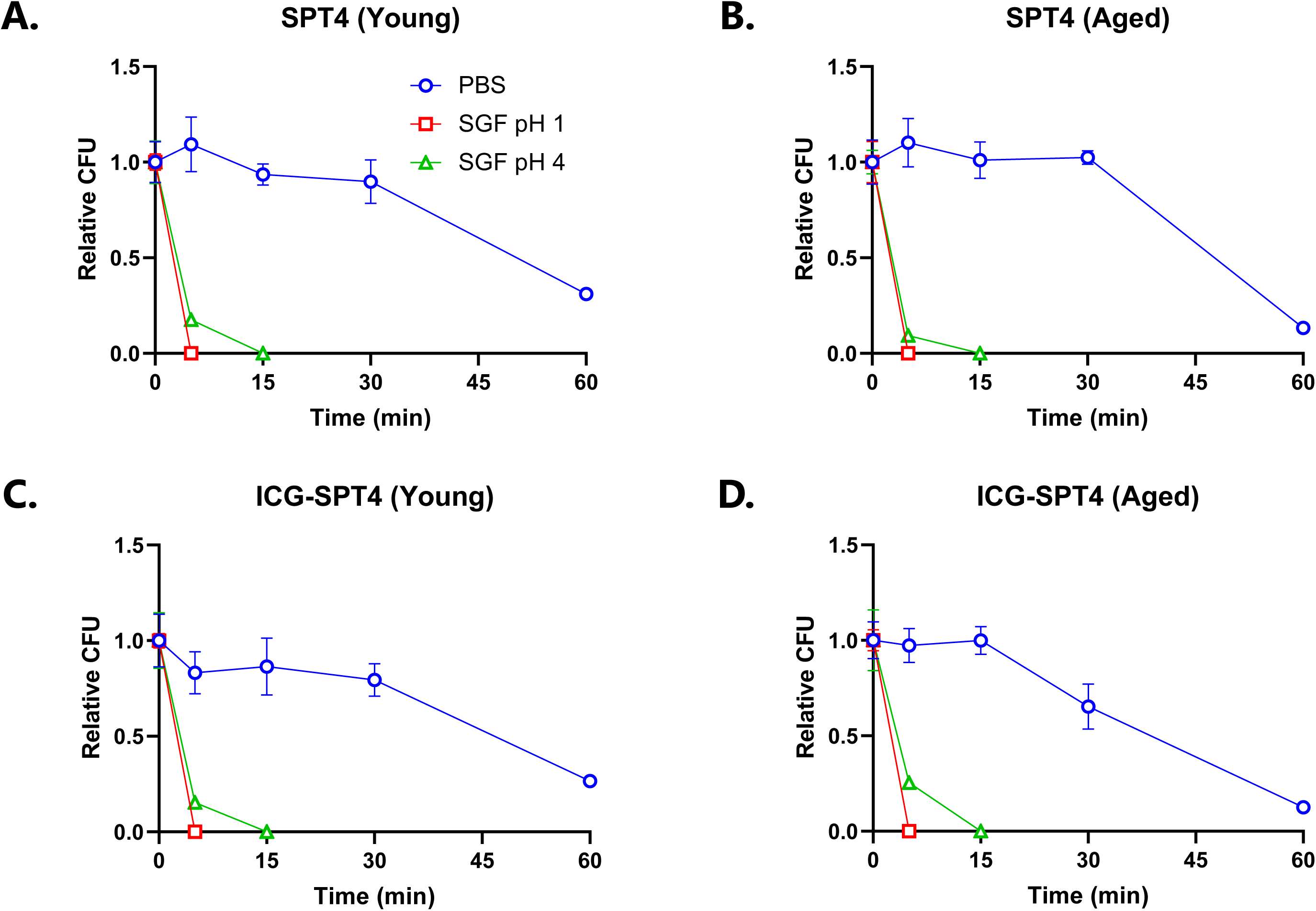
Susceptibility of SPT4 to SGF. Relative CFU of SPT4 (A, B) and ICG–SPT4 (C, D) prepared from young-mouse (A, C) and aged-mouse (B, D) inocula following exposure to SGF at pH 1.0–1.2 (SGF pH 1; fasting conditions) or 3.8–4.1 (SGF pH 4; postprandial conditions), with exposure to PBS as a control. CFU was measured at 5, 15, 30, and 60 min and normalized to the mean value of the corresponding group at time 0. Values below the detection limit are shown as 0. Data are presented as mean ± standard deviation (n = 3 per time point).

### Early aspiration in aged mice activates inflammatory and hematopoietic programs

To characterize early transcriptional responses, bulk RNA-seq was performed on cardiac blood collected 24 h post-inoculation (n = 2 per group), and DEGs were identified for the two primary comparisons (Aged_Asp vs. Aged_Col and Aged_Asp vs. Young_Asp; P < 0.05 and |log_2_FC| > 1). In the Aged_Asp vs. Aged_Col comparison, 374 genes were upregulated, and only 16 were downregulated in Aged_Asp mice (Figure 5A). In the Aged_Asp vs. Young_Asp comparison, 254 genes were upregulated, and 173 were downregulated in Aged_Asp mice (Figure 5A).

**Figure 5.**
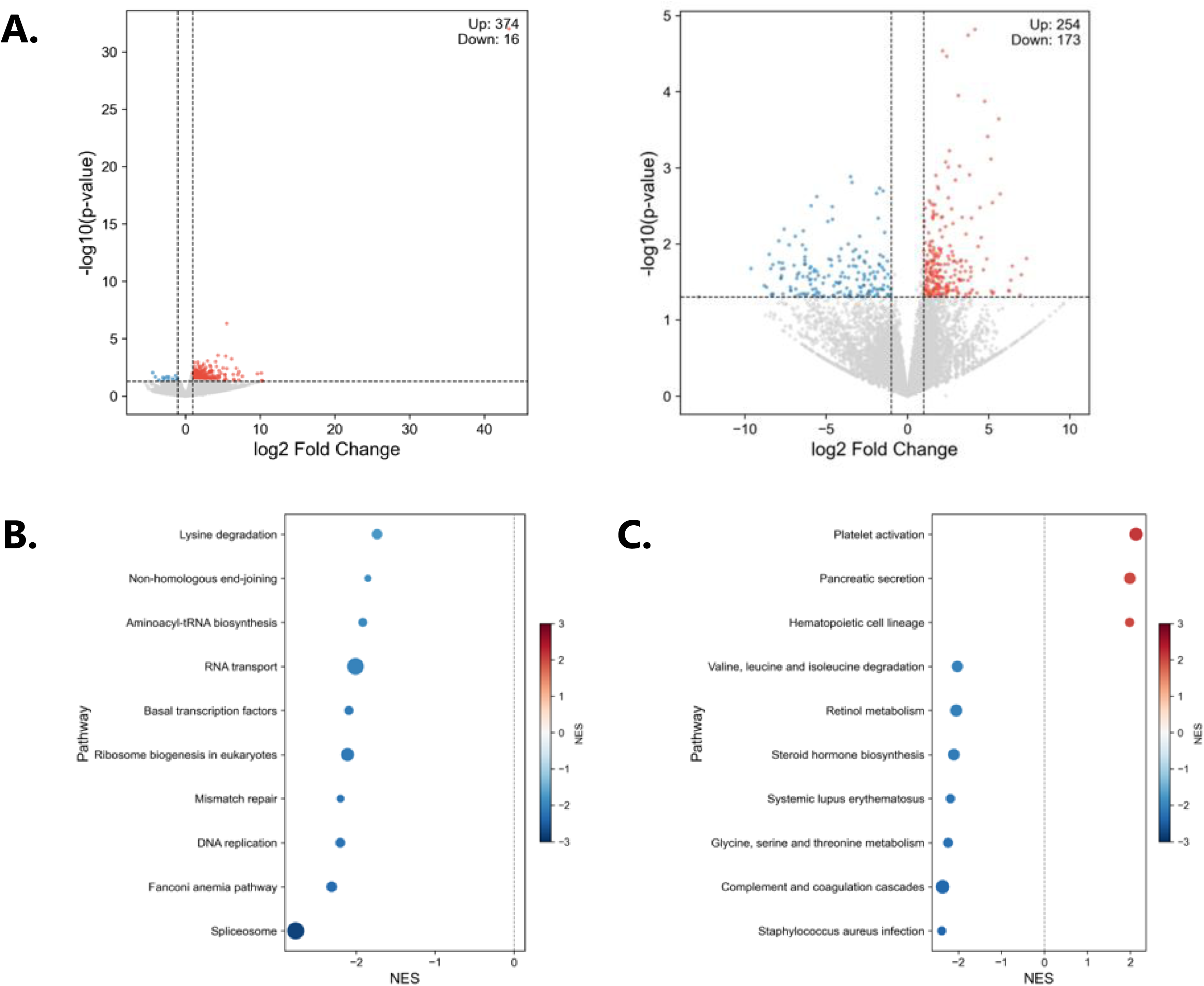
Blood transcriptomic profiles at 24 h post-inoculation. (A) Volcano plots showing DEGs (P < 0.05 and |log_2_FC| > 1) in Aged_Asp vs. Aged_Col (left) and Aged_Asp vs. Young_Asp (right) comparisons. Aged_Asp is the test group in both comparisons; red and blue points indicate upregulated and downregulated genes, respectively, in Aged_Asp mice. (B, C) GSEA of KEGG pathways for Aged_Asp vs. Aged_Col (B) and Aged_Asp vs. Young_Asp (C) comparisons. Dot color indicates the normalized enrichment score, and dot size indicates the gene-set size; n = 2 per group.

GSEA of KEGG pathways was performed for the two comparisons (Figure 5B, C). In the Aged_Asp vs. Aged_Col comparison, biosynthetic and proliferative pathways were negatively enriched in Aged_Asp mice, whereas inflammatory mediators, such as Il10, Saa2, and Arg1, were upregulated. In the Aged_Asp vs. Young_Asp comparison, platelet activation and hematopoietic cell-lineage pathways were positively enriched in Aged_Asp mice. These findings indicated that early aspiration in aged mice activated inflammatory and hematopoietic programs by 24 h after inoculation.

### Healthy aging reduces plasma innate immune and phagocytic proteins without an overt acute-phase response

To establish an age-related baseline in the absence of infection, we compared the plasma proteomes of healthy young (2-month-old; n = 6) and aged (18-month-old; n = 3) mice by data-independent-acquisition LC–MS/MS. Differential expression analysis identified 204 differentially expressed proteins (DEPs), of which 47 were increased and 157 were decreased in aged relative to young mice (Figure S1A, Data Set S1).

Notably, the healthy aged plasma proteome did not show an overt elevation of classical acute-phase reactants: serum amyloid A (SAA1) was in fact significantly lower in aged mice, and C-reactive protein, complement C3, haptoglobin, lipocalin-2 and S100A8/A9 were unchanged. Instead, several mediators of innate immune responsiveness were reduced in aged controls, including the Toll-like-receptor adaptor MYD88 and the interferon-inducible GTPase GBP6. As an exploratory analysis, we performed over-representation analysis using both the KEGG and Gene Ontology biological process (GO BP) databases (Figure S1B–E). KEGG returned few terms (ribosome among downregulated proteins; nitrogen metabolism, synaptic vesicle cycle and arginine biosynthesis among upregulated proteins), whereas GO BP highlighted a reduction in phagocytosis and endocytosis among downregulated proteins, with upregulated proteins dominated by neuronal and metabolic terms rather than inflammatory pathways. Given the small aged control group (n = 3), these enrichment results are presented as exploratory.

These findings indicate that, already before infection, aged mice exhibit a reduced capacity to mount innate immune and phagocytic responses.

### Early aspiration in aged mice engages complement, coagulation, and phagocytic pathways

PCA of all quantified proteins after filtering and imputation showed no clear group-level separation but substantial inter-individual variability, consistent with heterogeneous early host responses 24 h post-inoculation (Figure S2A). PC1 and PC2 accounted for 20.3% and 11.4% of the total variance, respectively.

Pairwise differential abundance analysis revealed that in the Aged_Asp vs. Aged_Col comparison, 69 proteins were upregulated, and 75 were downregulated in Aged_Asp mice (Figure 6A, Data Set S1). In the Aged_Asp vs. Young_Asp comparison, 118 proteins were upregulated, and 203 were downregulated in Aged_Asp mice (Figure 6A, Data Set S1).

**Figure 6.**
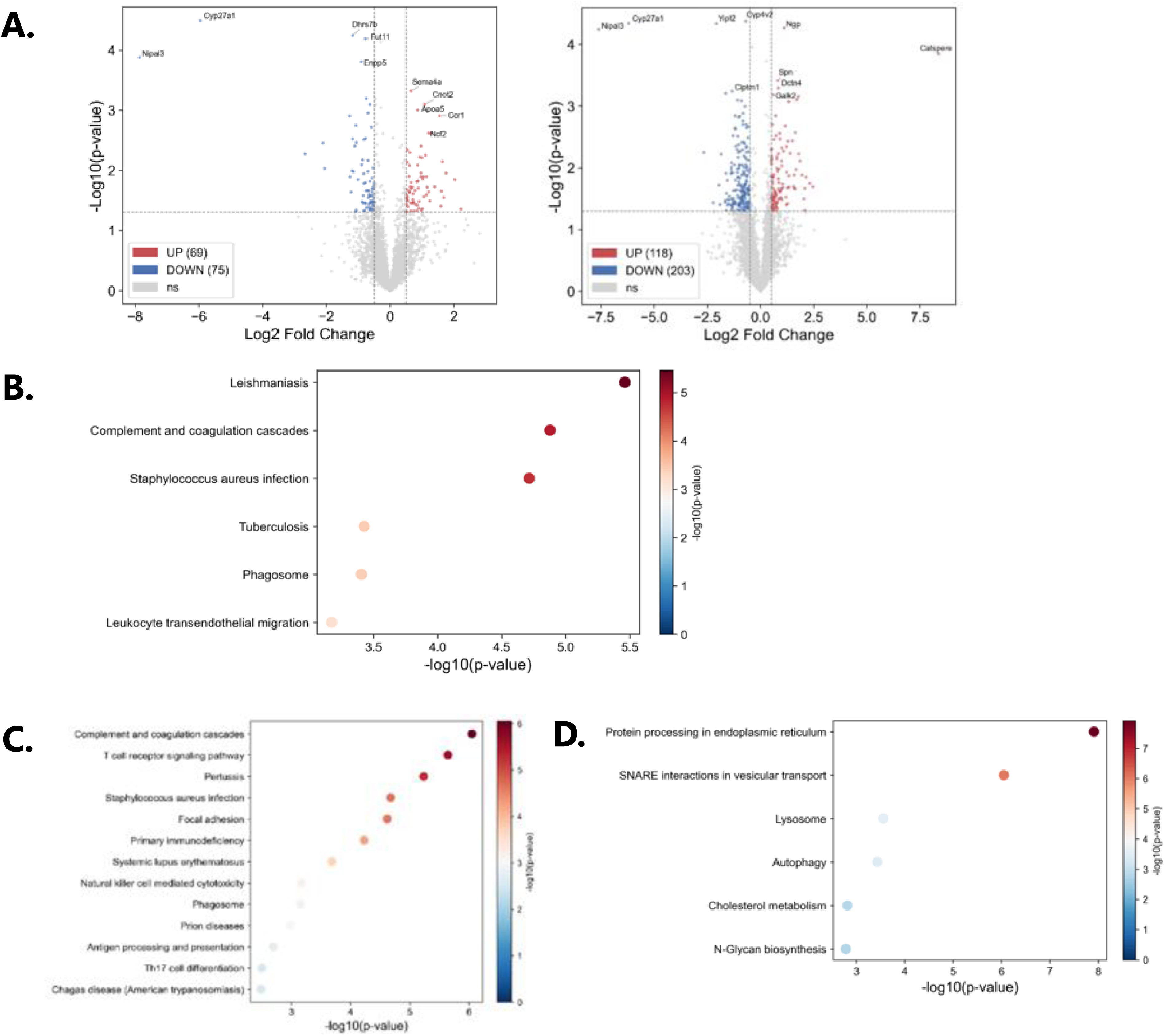
Plasma proteomic profiles at 24 h post-inoculation. (A) Volcano plots showing DEPs (P < 0.05 and |log_2_FC| > 0.5) in Aged_Asp vs. Aged_Col (left) and Aged_Asp vs. Young_Asp (right) comparisons. Aged_Asp is the test group in both comparisons; red and blue points indicate upregulated and downregulated proteins, respectively, in Aged_Asp mice. (B–D) Over-representation analysis of KEGG pathways for proteins upregulated in Aged_Asp vs. Aged_Col (B), upregulated in Aged_Asp vs. Young_Asp (C), and downregulated in Aged_Asp vs. Young_Asp (D) comparisons. PCA and PPI networks are provided in Figure S2. Young_Asp, n = 4; Aged_Asp, n = 4; Aged_Col, n = 4.

### Interaction network analysis of the plasma proteome resolves complement- and myeloid-associated modules

Protein–protein interaction network analysis was performed separately for proteins increased and decreased in Aged_Asp mice in the two primary comparisons (Figure S2B, C). In the Aged_Asp vs. Young_Asp comparison, increased proteins formed a densely connected network including complement- and immune-associated proteins such as C1QA, C1QB, C1QC, CTSS, STAT3, and VWF, whereas decreased proteins formed a separate network containing proteins such as CANX, LAMP1, RPN1, PDIA3, and STK11. In the Aged_Asp vs. Aged_Col comparison, increased proteins were organized around myeloid- and inflammatory response-related nodes including ITGAM, FCGR3, CCR1, C5AR1, NCF2, and NCF4, while decreased proteins formed smaller interaction networks including TGOLN1, TXNDC5, TSPAN2, and YIF1B.

Over-representation analysis of KEGG pathways was performed for both comparisons. Proteins upregulated in Aged_Asp mice were enriched predominantly for immune-related pathways, most notably complement and coagulation cascades and phagocytic pathways, in both Aged_Asp vs. Aged_Col (Figure 6B) and Aged_Asp vs. Young_Asp (Figure 6C) comparisons. In the Aged_Asp vs. Young_Asp comparison, proteins downregulated in Aged_Asp mice were enriched for pathways related to intracellular protein processing, vesicular transport, and metabolism (Figure 6D); no pathways were significantly enriched among proteins downregulated in the Aged_Asp vs. Aged_Col comparison. Collectively, proteins upregulated with aspiration mapped to complement, coagulation, and phagocytic pathways.

### Early aspiration in aged mice elicits transcriptomic and proteomic responses with limited molecular-level overlap

To identify molecular changes concordant at both transcript and protein levels, we integrated RNA-seq and plasma proteomic datasets for the two primary comparisons by matching genes and their encoded proteins. For each comparison, genes were classified as significant in the transcriptome (*P* < 0.05 and |log_2_FC| > 1), proteome (*P* < 0.05 and |log_2_FC| > 0.5), or both layers. Only a small number of features reached significance in both layers: 4 in the Aged_Asp vs. Aged_Col comparison (Figure 7A) and 13 in the Aged_Asp vs. Young_Asp comparison (Figure 7B). To assess concordance beyond these overlapping features, we compared transcript- and protein-level log_2_FC values across all 3,887 genes detected in both datasets (Figure 7C, D). The two layers showed only weak positive correlations, which were significant in the Aged_Asp vs. Aged_Col comparison (*r* = 0.053, *P* = 9.16 × 10 ⁴) but not in the Aged_Asp vs. Young_Asp comparison (*r* = 0.031, *P* = 0.051).

**Figure 7.**
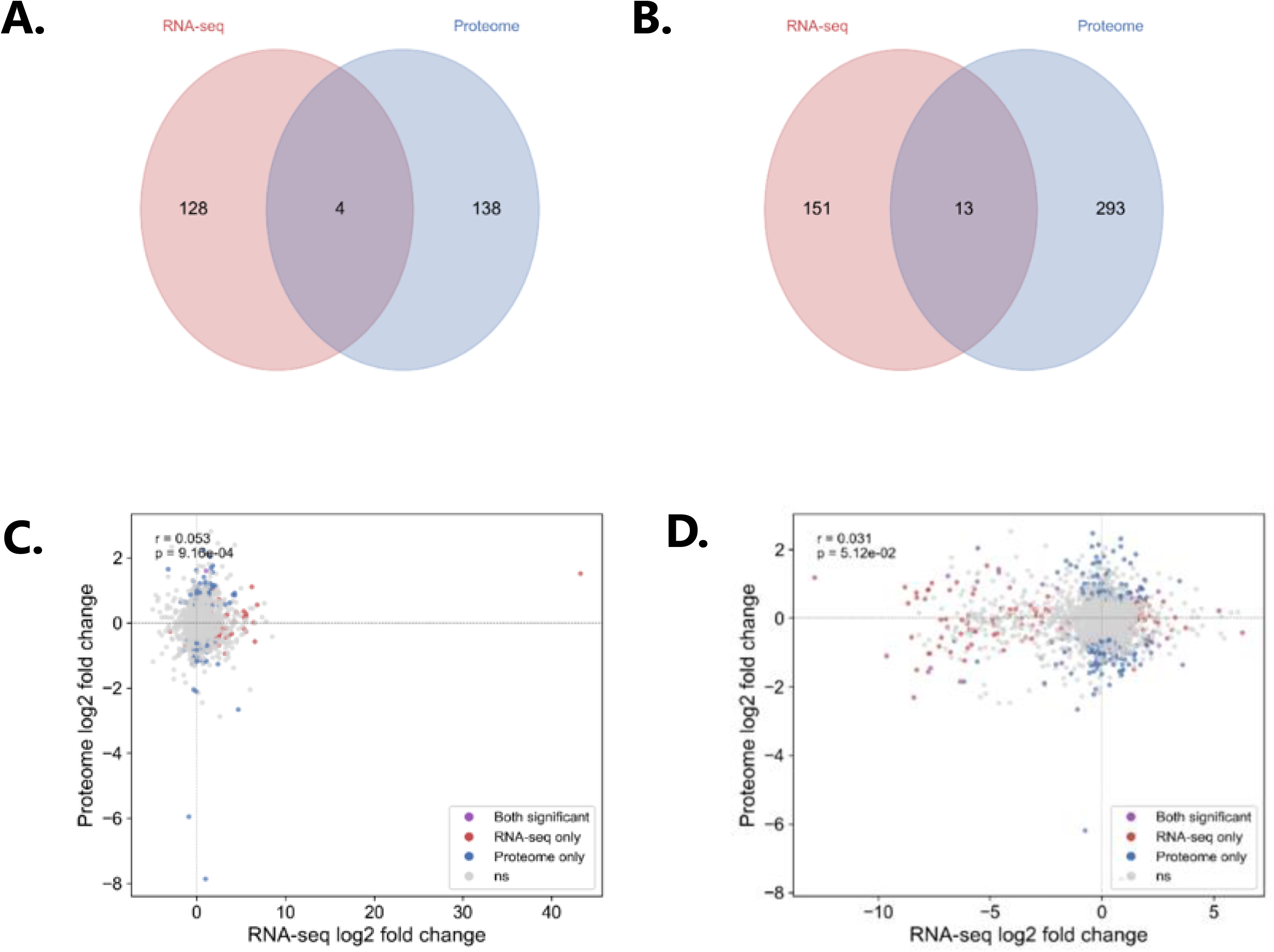
Multi-omics integration of RNA-seq and plasma proteomics. (A, B) Venn diagrams showing the overlap between significant DEGs (|log_2_FC| > 1, P < 0.05) and DEPs (|log_2_FC| > 0.5, P < 0.05) in Aged_Asp vs. Aged_Col (A) and Aged_Asp vs. Young_Asp (B) comparisons. (C, D) Scatter plots comparing RNA-seq and proteome log_2_FC for all genes detected in both datasets in Aged_Asp vs. Aged_Col (C) and Aged_Asp vs. Young_Asp (D) comparisons. Points are colored by significance: both significant (purple), RNA-seq only (red), proteome only (blue), and not significant (gray). Pearson correlation coefficients and P-values are shown in each panel.

## Discussion

Early pneumococcal infection is a temporally dynamic process, in which neutrophil recruitment, cytokine release, and monocyte mobilization evolve rapidly and asynchronously among individuals. In our model, mortality did not begin until day 3; therefore, the 24-h sampling window preceded the onset of death, positioning our profiling as an assessment of the early inflammatory response that may prefigure divergence in survival outcomes. Within this window, we used real-time NIR fluorescence imaging of ICG–SPT4 to stratify mice into colonization and aspiration groups during the first 10 min of intranasal inoculation. Survival analysis demonstrated that aspiration into the lower respiratory tract was associated with significantly higher mortality in both young and aged mice, indicating that the initial site of bacterial deposition was a key determinant of disease outcome.

Integration of transcriptomic and proteomic datasets revealed limited concordance at the level of individual features, with a few genes and proteins reaching significance in both layers and weak fold-change correlations. However, at the pathway level, the two datasets converged: proteins upregulated in the aspiration groups were consistently enriched for complement and coagulation cascades and phagocytic, myeloid-associated innate immune pathways, paralleling the inflammatory and hematopoietic programs prominent in the transcriptome. We regard this pathway-level convergence across two independent molecular layers as a robust readout of the early aspiration response. The limited feature-level overlap is expected at this early time point. The blood-cell transcriptome and plasma proteome sample distinct compartments and molecular pools with differing kinetics, and pronounced inter-individual variability is intrinsic to this early inflammatory phase. Whereas procedural variability was minimized by prospective, imaging-based stratification, this biological variability is an accepted feature of the model, rather than a technical artifact; our interpretation, therefore, rests on the convergent pathways rather than individual features.

This decline in innate immune responsiveness and macrophage/phagocyte function is a well-documented feature of immunosenescence and inflammaging [9], and it provides a plausible baseline explanation for the greater vulnerability of aged mice to aspiration-associated pneumococcal infection observed here. Notably, the “low-grade chronic inflammation” component of inflammaging was not accompanied by a pronounced systemic acute-phase response at the plasma level in this model, indicating that the age-associated predisposition in this model resides in diminished innate effector capacity rather than in a heightened basal inflammatory tone.

A key methodological consideration is the definition of the aspiration group based on NIR fluorescence signal detected in the lower respiratory tract. Although we did not directly confirm bacterial deposition in the lung parenchyma by bronchoalveolar lavage or CFU quantification, the anatomy of the rodent upper airway supported the interpretation that signals detected in the lower respiratory tract resulted from aspiration. Rodents possess an intranarial larynx, in which the epiglottis is positioned high against the soft palate so that the laryngeal inlet opens directly into the nasopharynx [48]. In the awake animal, this configuration favors airway protection: material entering the pharynx is directed laterally to the esophagus, and swallowing is accompanied by brief laryngeal closure and cessation of breathing that normally prevent aspiration [49]. Under general anesthesia, however, these protective swallowing and laryngeal-closure reflexes are suppressed, so that a liquid bolus delivered to the nasopharynx can enter the directly apposed laryngeal inlet and be aspirated into the lower airway rather than cleared to the esophagus. This anesthesia-dependent susceptibility is the basis of established oropharyngeal aspiration models used for pulmonary delivery in rodents [50], and is consistent with our observation that only a subset of anesthetized mice developed lower respiratory-tract signal. Accordingly, fluorescence extending beyond the nasopharyngeal region was prospectively verified as aspiration in each animal by real-time imaging.

Consistent with this interpretation, the same colonization- and aspiration-type signal distributions were reproducible on a conventional IVIS optical imager (Figure S3), indicating that the two patterns were not an artifact of our benchtop system but detectable across independent imaging platforms. Nevertheless, because the aspiration signal was transient, single-time-point still imaging on this platform could not reliably classify the two patterns, underscoring the value of continuous real-time acquisition for prospective, per-animal verification.

The survival advantage of the colonization group raises the question whether nasopharynx-confined bacteria pose systemic risk through alternative routes such as gastrointestinal translocation. NIR fluorescence imaging revealed that in some colonization mice, fluorescence signals were transiently detected in the gastric region, suggesting that a proportion of inoculated bacteria might have been swallowed rather than aspirated (Figure S4). However, SGF assay showed that SPT4 was rapidly killed under both fasting and postprandial gastric conditions, indicating that swallowed pneumococci were unlikely to survive the gastric transit. Because gastrointestinal translocation is dependent on species and host condition [51], the acid sensitivity of S. pneumoniae further underscores direct aspiration into the lower respiratory tract as the critical route of infection in this model.

S. pneumoniae is a natural pathogen of humans but not of rodents, and species-specific mismatches in major virulence factors contribute to relative resistance of mice to spontaneous pneumococcal infection of the lower respiratory tract. Consequently, experimental modeling generally requires high inocula, mouse-virulent strains, and deliberate delivery strategies. Each strategy comes with distinct advantages and limitations, rather than representing a clear hierarchy of validity. Conventional models of nasopharyngeal colonization rely on spontaneous transition of S. pneumoniae from the upper to lower respiratory tract, a physiological process, which is infrequent and difficult to time [12, 16]. Influenza A virus co-infection has been used to promote this transition [16, 52]; however, it introduces virus-associated inflammation as a confounding variable when the objective is to study pneumococcal response in isolation. Direct intratracheal or oropharyngeal instillation delivers a controlled dose reproducibly without viral co-infection [50]; however, it bypasses the upper airway and, therefore, neither reproduces the physiological branch point nor allows the initial site of deposition to be examined as a determinant of outcome.

The species specificity noted above is evident at the level of individual virulence factors, several of which engage host receptors in a human-restricted manner. The surface adhesin choline-binding protein A (CbpA; also known as PspC, SpsA, or Hic) binds the human polymeric immunoglobulin receptor (pIgR) and secretory component to mediate adherence to and translocation across the nasopharyngeal epithelium, yet it does not recognize the corresponding proteins of mice, rats, or rabbits [53, 54]. Similarly, CbpA recruits the complement alternative-pathway regulator factor H to evade opsonophagocytosis, but this interaction is restricted to human factor H and does not occur with the murine protein; accordingly, deletion of the factor H-binding domain of CbpA does not attenuate virulence in a mouse bacteremia model [55, 56]. A further species difference involves the phosphorylcholine-dependent engagement of the platelet-activating factor receptor (PAFR) that is required for epithelial invasion [12, 13].

In the present study, we took an alternative approach by exploiting the anatomical susceptibility of anesthetized mice to aspiration, enabling direct and prospectively verifiable delivery of bacteria to the lower respiratory tract in the absence of viral co-infection. Unlike intratracheal instillation, real-time imaging prospectively identifies animals with lower respiratory tract deposition before the onset of severe disease. Because this design preserves the physiological branch point between colonization and aspiration, the initial site of bacterial deposition can be examined as a determinant of outcome, making the model well suited for studying aspiration-associated pneumonia in aged hosts, as directly addressed by the present young vs. aged comparison. ICG-based stratification ensures that only mice with confirmed lower respiratory tract deposition are included in the aspiration group, so that the systemic molecular signatures observed in this group can be attributed to bacterial aspiration itself, rather than viral co-infection or retrospective misclassification. Beyond these delivery considerations, the labeling chemistry itself offers practical advantages. In contrast to metabolic labeling approaches that incorporate probes into the bacterial cell wall, our method relies solely on simple mixing and centrifugation, requiring no genetic modification or chemical conjugation and providing a rapid, broadly applicable approach well suited to the short-term, binary readout used here.

The colonization group plays a dual role. For the mechanistic question addressed here, the contrast between colonization and aspiration is essential; in intervention studies evaluating vaccines or therapeutics, prospective enrichment for confirmed aspiration may instead be advantageous for animal reduction

Because colonized animals largely survive without progressing to lethal disease, their inclusion in conventional infection models that cannot resolve the site of deposition dilutes any treatment effect and inflates the group sizes required to reach statistical significance. Prospective enrichment for confirmed aspiration removes this source of variability and should therefore lower the number of animals needed to detect efficacy. A remaining consideration is that, under the current protocol, aspiration occurs in only a subset of inoculated animals, so that obtaining a target number of aspiration animals requires inoculating more animals than ultimately contribute evaluable data. Further refinement of the inoculation procedure to raise the proportion of animals that aspirate would be a welcome addition in this regard, though the imaging-based enrichment already provides the principal benefit for animal reduction in intervention studies.

Taken together, our findings establish real-time NIR fluorescence imaging as a means of prospectively resolving the initial site of pneumococcal deposition, and demonstrate that aspiration into the lower respiratory tract, rather than nasopharyngeal colonization, triggers severe disease across host ages. Multi-omics profiling at 24 h linked this outcome to an early host response, in which inflammatory and hematopoietic transcriptional programs and complement, coagulation, and phagocytic proteomic pathways were concurrently engaged. By preserving the physiological branch point between colonization and aspiration, this model provides a tractable platform for dissecting aspiration-associated pneumonia in aged hosts, and for evaluating preventive or therapeutic interventions in a clinically relevant setting.

This study has some limitations. First, the cohorts were modest, and the multi-omics groups were particularly small (notably the young colonization group, n = 2); therefore, the omics comparisons were exploratory, defined using nominal P-values without multiple-testing correction, with interpretation resting on pathway-level convergence rather than individual features (Figure 7). Second, profiling was performed at a single early time point (24 h) and did not resolve the kinetics of host response. Third, only male mice at two ages (2 and 18 months) were studied, leaving sex-specific effects and the full aging trajectory unaddressed. Fourth, aspiration was assigned from lower-respiratory-tract NIR signal rather than direct confirmation by bronchoalveolar lavage or CFU. Finally, because S. pneumoniae is a natural pathogen of humans but not rodents, findings should be extrapolated to human disease with caution. In the future, large, longitudinal cohorts of both sexes with direct confirmation of pulmonary bacterial delivery will be needed to build on these findings.

## Acknowledgements

We thank Mayumi Ishida for technical assistance with the proteomics experiments. This study was partly completed using SQUID at the D3 Center, University of Osaka, Japan, under the ’Joint Usage/Research Center for Interdisciplinary Large-scale Information Infrastructures (JHPCN)’ (Project ID: jh250016 and jh260013). Aged C57BL/6J mice were provided by the RIKEN BRC through the National BioResource Project (NBRP) of MEXT/AMED, Japan. We acknowledge the NGS core facility at Research Institute for Microbial Diseases of The University of Osaka for sequencing.

## Funding

This work was supported by JSPS KAKENHI Grant Numbers 23K27764, 25K13665, and 26K23623, and the Joint Research Program of the Research Center for GLOBAL and LOCAL Infectious Diseases, Oita University (2025B01).

## Disclosure statement

The authors report there are no competing interests to declare. Declaration of generative AI use

During the preparation of this manuscript, the authors used a generative AI tool (Claude, Anthropic; Claude Opus 5) to assist with language refinement, formatting of the reference list, organization of data outputs, and troubleshooting and debugging of data-analysis code.

## Data availability statement

The datasets generated and analyzed in this study have been deposited in public repositories and are accessible to the editors and reviewers for the purposes of peer review. The bulk RNA-seq data have been deposited in the NCBI Gene Expression Omnibus (GEO) under accession number GSE342088. The data can be accessed at https://www.ncbi.nlm.nih.gov/geo/query/acc.cgi?acc=GSE342088 using reviewer token ofinscgstvejxwf. The mass spectrometry proteomics data have been deposited to the ProteomeXchange Consortium via the jPOST repository with the dataset identifier PXD061351. The data can be accessed at https://repository.jpostdb.org/preview/11440689276a6aaf8439f08 using the access key 5179.

Both omics datasets are currently under confidential pre-publication access, which is the standard deposition state offered by these repositories prior to article publication. This arrangement was agreed in advance among the participating research groups, as the sequencing and proteomic analyses were performed as a collaboration. Both records are scheduled for full public release, with no further restriction and without any requirement for login credentials, upon acceptance of this article and in any case no later than its date of publication.

The source data underlying all figures, comprising the individual values behind every reported mean and standard deviation, the values used to construct each graph, and the quantified values extracted from images for analysis, are openly available without embargo and without any requirement for login credentials in figshare under the digital object identifier https://doi.org/10.6084/m9.figshare.33457540, released under a Creative Commons Attribution 4.0 International (CC BY 4.0) licence. This record also contains the differentially abundant protein lists cited in the text as Data Set S1.

Supplementary Videos 1 and 2, showing representative real-time near-infrared recordings of the colonization and aspiration patterns, are openly available in the same repository record under the digital object identifier https://doi.org/10.6084/m9.figshare.33457540 (CC BY 4.0).

No data underlying this study have been withheld from the editors or reviewers. All experiments were performed in mice; the study involved no human participants and generated no personally identifiable or otherwise sensitive data, and no ethical, privacy, or security restriction applies to the release of any dataset described above.

## Author contributions statement

T.S., Conceptualization, Methodology, Investigation, Formal analysis, Data curation, Visualization, Funding acquisition, Writing - original draft, Project administration; M.K., Methodology, Investigation; Z.S., Investigation, Validation; S.M., Investigation, Resources, Writing - review and editing; D.M., Investigation, Resources; T.Y., Investigation; S.S., Investigation; J.A., Conceptualization, Methodology, Supervision, Writing - review and editing; M.Y., Conceptualization, Supervision, Funding acquisition, Writing - review and editing.

## Notes

### Competing Interest Statement

The authors have declared no competing interest.

### Summary of Updates

The manuscript has been reformatted for a different target journal. As part of this reformatting, material previously presented as supplementary results has been moved into the main text. The main text is therefore substantially longer than in the previous version, although no new data were acquired and no findings were changed. Methods expanded for ARRIVE 2.0 compliance: study periods, animal sources, housing and husbandry conditions, anesthesia and reversal agent doses, imaging system parameters, and mass spectrometry acquisition settings are now reported in full. Results section revised for accuracy: the description of principal component analysis of the plasma proteome was consolidated, and the statement of statistical comparisons for bacterial growth characteristics was restricted to the labeled versus unlabeled comparisons actually tested. Discussion revised to remove redundant text. The comparative anatomy of the rodent upper airway is now described in one place rather than two, and duplicated statements on host species specificity of pneumococcal virulence factors were removed. A reference to a supplementary discussion section was removed, as that material has been incorporated into the main text. Figure captions revised: Figure 1 now specifies the scope of the statistical comparisons performed; Figure 4 now states the normalization procedure and notes that values below the detection limit are plotted as zero. Reference list reordered to follow order of first citation in the revised text; references are renumbered accordingly. Data availability statement updated. Source data for all figures, the differentially abundant protein lists, and the supplementary videos are now openly available in figshare under DOI 10.6084/m9.figshare.33457540 (CC BY 4.0), with no embargo and no login requirement. Supplemental files updated: the Supporting Information now includes the supplementary data set of differentially abundant proteins, and the supplementary video captions were made consistent with the main text. Typographical corrections were applied throughout, including consistent use of en dashes in compound terms, superscript and subscript formatting for units and variables, and consistent abbreviation definitions at first use. No new experiments were performed, and no data, statistical results, or conclusions were changed in this revision.

