## Supplemental Figures for "Real-time near-infrared imaging distinguishes nasopharyngeal colonization from aspiration of *Streptococcus pneumoniae* and identifies aspiration as a trigger of severe disease"

**Supporting Information (SI)**

**(A)**

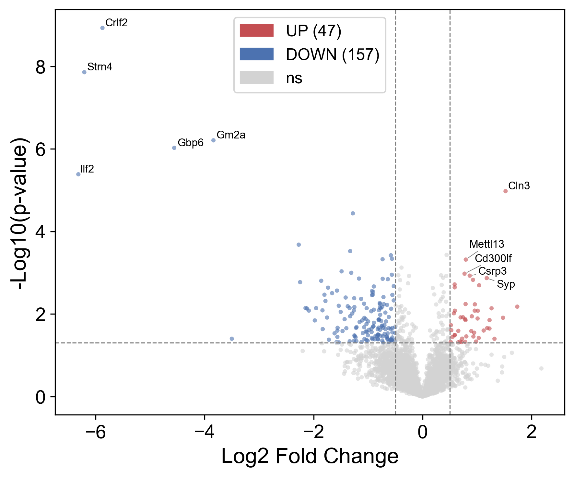

**(B) UP	(C) DOWN**

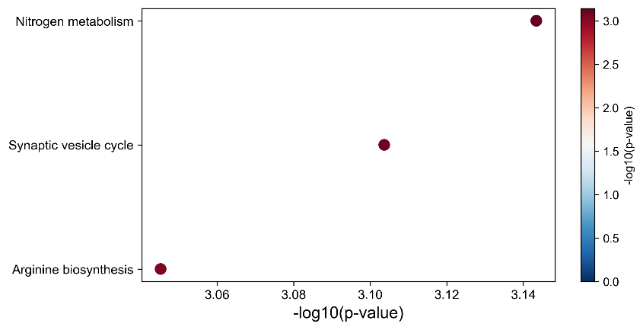

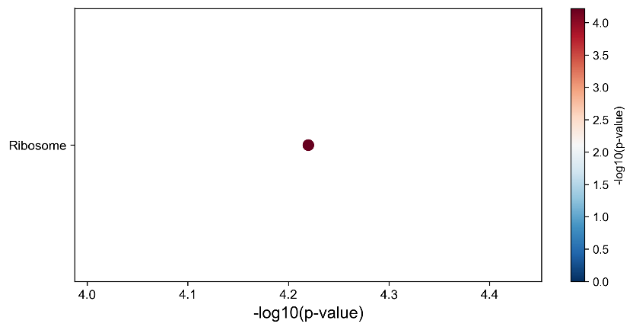

**(D) UP**

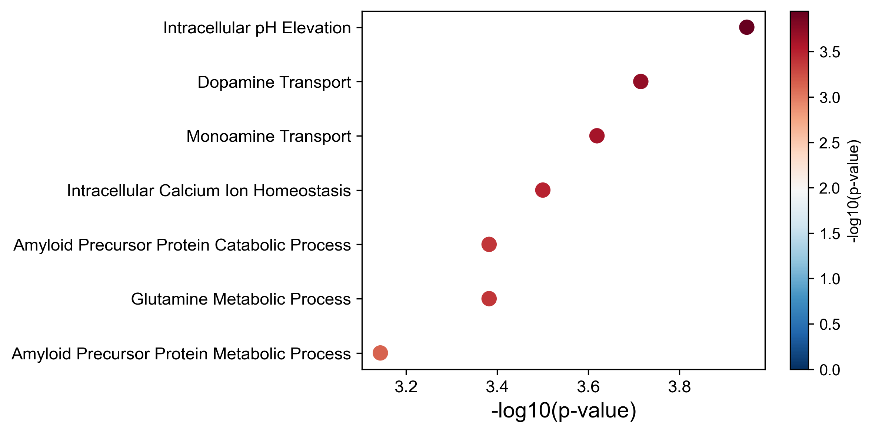

**(E) DOWN**

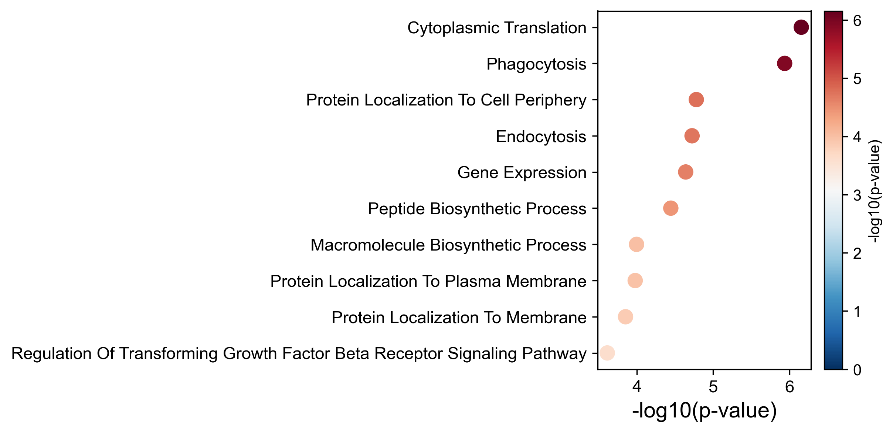

**Figure S1. Plasma proteomic signature of healthy aging (young vs. aged control mice).** (A) Volcano plot of aged vs. young control mice (test = aged, reference = young); red, increased in aged (47 proteins); blue, decreased in aged (157 proteins); dashed lines indicate |log₂FC| = 0.5 and *P* = 0.05. (B, C) Exploratory over-representation analysis (Enrichr, KEGG) of proteins (B) upregulated and (C) downregulated in aged mice. (D, E) The same analysis using GO biological process, for proteins (D) upregulated and (E) downregulated in aged mice. Dot color and position indicate −log₁₀(*P). n =6 (young control) and n =3 (aged control).*

**(A)**

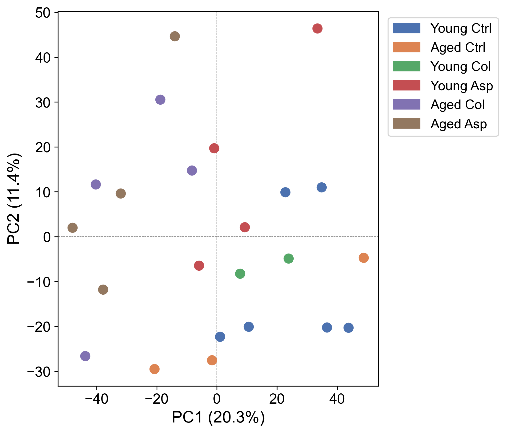

**(B) Aged_Asp vs. Young_Asp UP	Aged_Asp vs. Young_Asp DOWN**

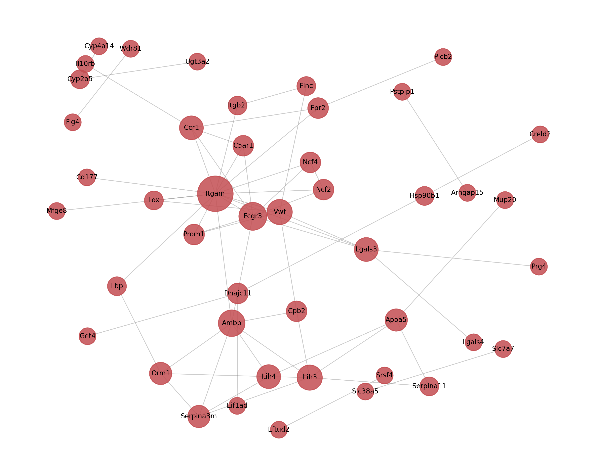

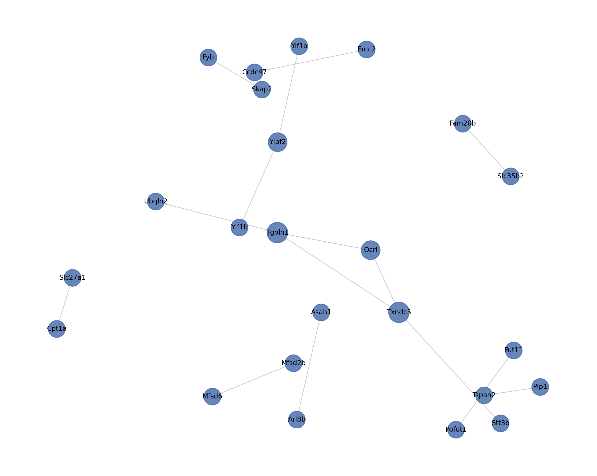

**(C) Aged_Asp vs. Aged_Col (Ref) UP	Aged_Asp vs. Aged_Col (Ref) DOWN**

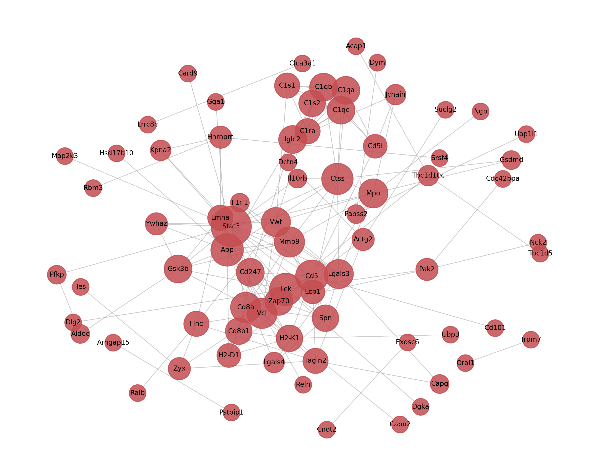

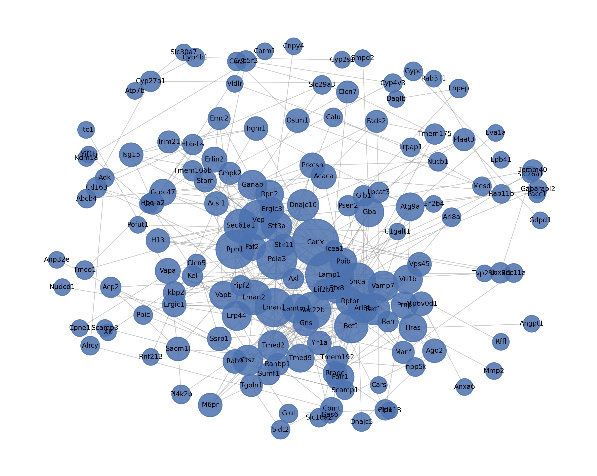

**Figure S2. Plasma proteomic profiles at 24 h post-inoculation.** (A) Principal component analysis of all quantified proteins after filtering and imputation; PC1 and PC2 account for 20.3% and 11.4% of the total variance, respectively. (B, C) Protein–protein interaction (PPI) networks of proteins increased (UP) and decreased (DOWN) in aged aspiration-type (Aged_Asp) mice, for the (B) Aged_Asp vs. Young_Asp and (C) Aged_Asp vs. Aged_Col comparisons.

**(A)**

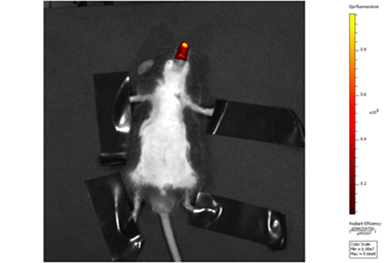

**(B)**

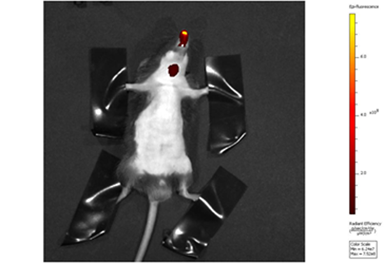

**(C)**

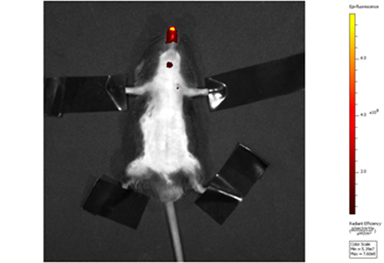

**Figure S3. *In vivo* NIR fluorescence imaging of ICG–SPT4 using a conventional IVIS system.** Mice were imaged with an IVIS optical imager immediately after the fourth intranasal inoculation. Representative images of a colonization-type mouse (A) and aspiration-type mice (B, C) are shown.

(A)	(B)

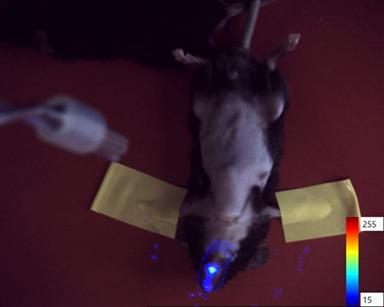

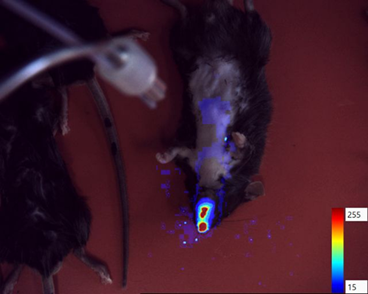

**Figure S4.** **Real-time NIR fluorescence imaging of ICG-labeled *S. pneumoniae* (ICG–SPT4) following intranasal inoculation.** Representative images of a young colonization-type (Col) mouse at 8 min (A) and 1 h (B) after intranasal inoculation. ICG–SPT4 was visualized as near-infrared (NIR) fluorescence overlaid on the visible-light image. In (B), a linear signal along the esophagus and a rounded signal in the underlying gastric region were also observed, consistent with swallowing of a proportion of the inoculum.

**Supplementary Videos**

**Supplementary Video 1. Real-time imaging of an aged aspiration-type (Aged_Asp) mouse during the first minutes after intranasal inoculation, shown as a merged visible-light and NIR fluorescence overlay.** ICG–SPT4 is tracked in real time as it distributes through the airway.

**Supplementary Video 2. The same acquisition as Supplementary Video 1, shown as the NIR fluorescence channel only, highlighting the distribution of ICG–SPT4.**

Supplementary Videos 1 and 2 are available in the figshare record (<https://doi.org/10.6084/m9.figshare.33457540>).
